# Evolving a High-Performance Terminal Deoxynucleotidyl Transferase for Enzymatic DNA Synthesis

**DOI:** 10.1101/2024.06.24.600205

**Authors:** Stephanie Forget, Mikayla Krawczyk, Anders Knight, Charlene Ching, Rachelle Copeland, Niusha Mahmoodi, Melissa Mayo, James Nguyen, Amanda Tan, Matt Miller, Jonathan Vroom, Stefan Lutz

## Abstract

Enzymatic DNA synthesis, using stepwise nucleotide addition catalyzed by template-independent polymerases, promises higher efficiency, quality, and sustainability than today’s industry standard phosphoramidite-based processes. We report on the directed evolution of a terminal deoxynucleotidyl transferase that uses 3’-phosphate blocked dNTPs to control the polymerization reaction and demonstrates high activity for these modified substrates and improved template promiscuity and thermostability.

## Main text

DNA synthesis is a cornerstone of biological research. From the assembly of new synthetic DNA constructs to the amplification or mutagenesis of existing sequences with oligonucleotide primers, research depends on the ability to write DNA accurately, quickly, and inexpensively. These constructs and primers are primarily synthesized using phosphoramidite-based chemistry first reported in 1981.^1^ Steady improvements and optimization of the process over the last four decades have driven up the average nucleotide coupling efficiency of this solid-phase synthesis to ≥99.5%.^2^ Despite such superb performance, the yield of full-length DNA falls off as oligonucleotide length exceeds 150-200 base pairs, and DNA quality is diminished by repeated exposure to harsh chemical conditions during the iterative synthesis. Furthermore, phosphoramidite synthesis has poor atom economy and generates significant hazardous organic waste.^3^ In contrast, enzymatic DNA synthesis can exploit the remarkable fidelity and efficiency of nature’s replication machinery while also offering a greener process that runs in aqueous conditions and minimizes the need for activators, oxidants, and protection groups.

Conceptually, enzymatic DNA synthesis of oligonucleotides is an iterative, two-step process of extension and deblocking. In the extension step, a polymerase catalyzes the template-independent addition of a 2’-deoxynucleoside triphosphate (dNTP) to the 3’-end of a polynucleotide strand. One strategy for limiting extension to a single nucleotide involves using polymerase-dNTP conjugates to sterically hinder the 3’ terminus of the extinction product.^4, 5^ Alternatively, run-away polymerization can be avoided via reversible blocking of the dNTP’s 3’-hydroxyl group, which gives the added benefit of producing scarless oligonucleotide products.^6^ In the deblocking step, hydrolysis liberates the 3’-position of the growing oligonucleotide and readies it for the next cycle of nucleotide addition. Critical for the translation of this concept into a competitive commercial DNA synthesis process are the high speed, coupling efficiency, and uniformity of each reaction step – goals most elegantly accomplished by an all-enzymatic approach.

Terminal deoxynucleotidyl transferases (TdTs) are the most well-studied polymerases in the context of template-independent synthesis. TdT was proposed as a catalyst for enzymatic DNA synthesis using nucleotides with blocked 3’-hydroxyl groups in 1962,^7^ and it has been a leading candidate for DNA synthesis applications ever since.^8-10^ Steric constraints in the TdT active site have in the past favored modifications with a small steric footprint such as nitrobenzyl^11^ or aminoalkoxyl^12, 13^ groups. However, removal of these entities requires photolytic or chemical unblocking/deprotection step which adds complexity to the overall process. Alternatively, a phosphate group is an effective blocking group^14^ that can efficiently be hydrolyzed by commercially available phosphatases under reaction conditions similar to the TdT-catalyzed nucleotide extension step.^15, 16^ These deblocking reactions can be carried out at physiological pH conditions that minimize depurination and other potential DNA damage. While 3’-phosphate blocked dNTPs (3’P-dNTPs) would enable an all-enzymatic solution to DNA synthesis, this large, polar blocking group is poorly tolerated by wild-type TdTs.

Protein engineering by directed evolution can be applied to address the functional impediment of TdT towards use of 3’P-dNTPs. At the same time, engineering efforts must select TdT variants with improvements to several traits required for high performance in DNA synthesis. Specifically, process requirements call for nucleotide incorporation efficiencies of >99% at reaction times of <90 seconds per cycle and tolerance of elevated reaction temperatures (60 °C) to minimize interference of 3’-terminal DNA secondary structure with TdT binding.^17^ In a departure from traditional enzyme engineering focused on tailoring one biocatalyst to one specific substrate,^18-20^ TdT evolution must strive for maximum substrate promiscuity. Native TdTs exhibit significant bias with respect to dNTPs and the three nucleotides at the 3’-terminus of the oligo acceptor sequence. Given 64 (4^3^) unique combinations of oligonucleotide 3’-terminal sequences and four 3□P-dNTP substrates, the ideal polymerase for enzymatic DNA synthesis must achieve high and uniform nucleotide coupling efficiency on 256 substrate pairs (Figure 1a). Herein, we describe the directed evolution of TdT for use in commercial enzymatic DNA synthesis.

**Figure 1.**
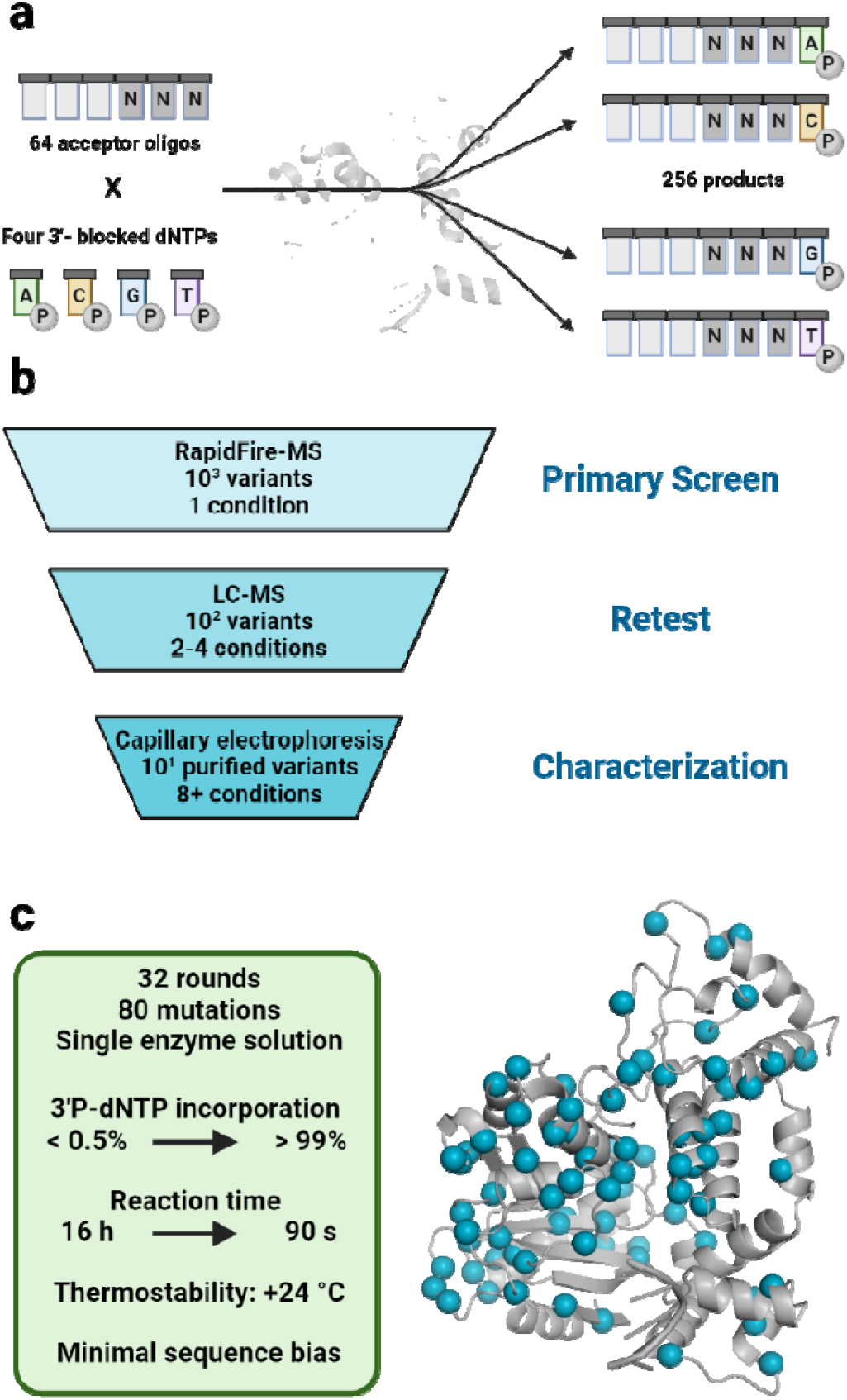
DNA synthesis problem and the enzyme engineering solution. (a) Combinatorial complexity of oligo synthesis with oligo sequence bias. Accounting for the three 3’-terminal nucleotides in an oligo acceptor and the four 3’P-dNTPs, an enzyme for DNA synthesis needs to catalyze 256 polymerization reactions with non-native substrates at a minimum of 99% coupling efficiency to be commercially relevant (b) Multiple tiered approach to screening. Thousands of variants can be screened in one day under a single reaction condition using a RapidFire-MS system. Variants (100-400) showing improvement under the primary screen are tested under multiple additional challenge conditions. Top variants from the secondary screening are shake-flask purified and characterized under several challenge conditions, with one variant selected as the backbone for the next round of evolution. (c) Mutations in **TdT-33** mapped onto a homology model. 80 mutations across 32 rounds of evolution brought the engineered **TdT-33** to high coupling efficiency and low sequence bias in 90-second reactions. Created with BioRender.com.

The *N*-terminally truncated TdT from *Empidonax traillii* was selected from a panel of wild-type TdTs as the evolution parent given its modest stability and heterologous expression in *E. coli*. This selected evolution parent (**TdT-01**) had trace starting activity with 3’P-dNTPs and required significant improvements to stability, solubility, and expression to become a viable commercial catalyst.

Evolution in the initial rounds focused on improving soluble expression and stability. To select for these traits, libraries were screened with a thermal challenge using 2’,3’-dideoxynucleotide triphosphates (ddNTP) as surrogate substrates. Heat-treated, clarified *E. coli* lysates were used as the source of TdT for screening. Crude *E. coli* lysates contain oligonucleotide-degrading nucleases and dNTP-degrading phosphatases, which potentially complicated analysis of screening results. However, mild heat treatment was found to be sufficient to reduce these side reactions and to allow for reliable monitoring of TdT product formation. Compared to the DNA substrate, ddNTP-extended products showed higher resistance to degradation under lysate screening conditions. It was therefore possible to monitor ddNTP-extended products using LC-MS to identify beneficial variants in the context of a crude lysate screen.

After three rounds of evolution, activity on the target 3’P-dNTP substrate was measurable but could not be easily screened in lysate due to interference from lysate components. However, improvements in the stability and solubility of TdT variants over the first rounds made it possible to recover sufficient yields of TdT using a high-throughput (HTP) purification protocol. In this protocol, cell pellets were resuspended in buffer containing lysozyme then subjected to a two-step purification procedure utilizing Ni-NTA affinity sollowed by desalting. Screening with the purified enzymes allowed for longer incubation times (1-2 hours) without product degradation, making it possible to screen for 3’P-dNTP incorporation despite low activity. At this point, HTP screening was being performed with relatively high-incorporating oligo substrates to find variants that would accommodate the 3’-phosphate blocking group.

Given the low starting activity with 3’P-dNTPs and initial evolution trajectory, we predicted that a significant number of rounds (>20) would be required to evolve TdT to meet target process requirements. Thus, we sought to simplify screening as much as possible to reduce operational complexity, time, and cost for each round of evolution. The HTP purification procedure that was initially required to screen with 3’P-dNTPs was lengthy and of limited throughput, so efforts were made to return to screening using lysates. As a result of cumulative activity and stability improvements, it became possible to screen reactions with 3’P-dNTPs using TdT in *E. coli* lysates by round 9 of evolution. Higher enzyme stability allowed for higher lysate thermal pre-treatments, which reduced degradation of oligo and 3’phosphate-dNTP by lysate components while improved activity enabled detection of 3’-phosphorylated oligo products with lower lysate loading and shorter reaction times. From this point forward, all HTP screening was with heat-treated cell lysates.

With these high-throughput lysate assays established, the subsequent rounds focused on increasing promiscuity, reducing bias towards oligo sequences, and improving incorporation efficiency for 3’P-dNTPs. In parallel, we continued to monitor and select for improvements in soluble expression and stability. By this point in evolution, an optimized workflow was developed (Figure 1b). Key to enabling the evolution was the development of fast, reliable, and robust analytical methods to enable tiered HTP screening at progressively process-relevant conditions.

Primary screening represents a first pass to identify active variants using a single reaction condition. The primary screening assay relied on a Rapid-Fire MS (Agilent) analytical method using a HILIC cartridge to swiftly analyze and select improved variants, which enabled screening of over 2000 variants overnight. Active and sequence unique variants identified in the primary screen were re-grown for additional testing with different screening pressures to improve reaction rate, thermal stability, or sequence bias. As part of this secondary screening, re-test plates (typically 100-200 variants) were screened under 2-4 additional challenge conditions to further vet individual mutations, and lower-throughput LC-MS assays were used to track additional species of interest. TdT amino-acid substitutions that produced byproducts such as such as multiply extended or 3’-dephosphorylated products were tracked and selected against when appropriate.

Mutations found to be beneficial in primary and re-test screening were folded into combinatorial libraries. Combinatorial libraries were evaluated by secondary screening, using 2-4 conditions and LC-MS analytical detection. A small set of variants (typically 6-8) with improved activities across multiple selection pressures were then screened in a third tier of validation assays using shake-flask-purified enzymes. These enzymes were evaluated for activity and stability improvements using either LC-MS or capillary electrophoresis (CE) such that a detailed analysis of the reaction profile could be obtained. Mirroring our approach to HTP screening, reactions with purified proteins were evaluated using screening pressures including short reaction times, low-activity substrates, and reduced 3’P-dNTP equivalents. In each round, the variant with the best overall improvement across multiple traits was selected as the parent for the next round of evolution. Throughout evolution, variants were screened against a panel of oligo substrates bearing variable sequences at the last three 2’-deoxyribonucleotide positions at the 3’-terminus, and under-performing substrates were identified for primary and re-test screening. Thirty-two rounds of evolution were completed using these approaches. Over the course of these rounds of evolution, each position in TdT (following the N-terminal His-tag) was targeted at least once, 75% of the single amino-acid substitution landscape was observed, and 80 mutations were incorporated into **TdT-33** with respect to **TdT-01** (Fig 1C). Representative TdT variants from the evolution lineage were selected for head-to-head comparison to illustrate progress (Figure 2).

**Figure 2.**
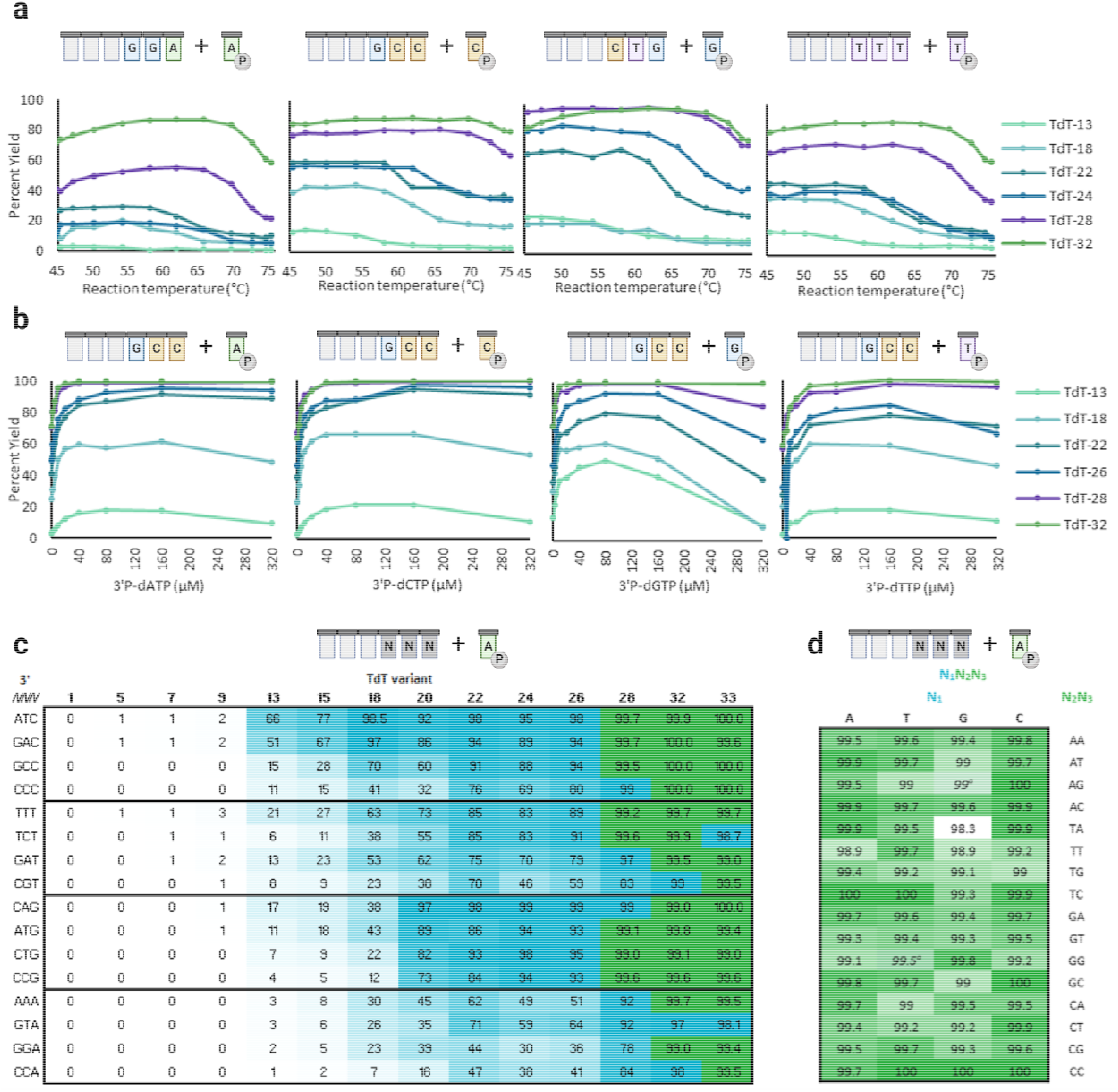
TdT performance improvements during directed evolution. A) TdT variant activity at varying temperatures. Residual TdT activity following a thermal challenge from 45 °C to 75 °C were assayed with each 3’P-dNTP and TdT variants from **TdT-13** to **TdT-32**. B) Reduced blocked nucleotide equivalent requirements through evolution. TdT activity was measured with varying concentrations of each 3’P-dNTP and TdT variants from **TdT-13** to **TdT-32**. C) Sequence bias improvements through evolution. Nucleotide coupling efficiency with 3’P-dATP and 16 varied oligonucleotide acceptors in a 90-second reaction using 14 TdT variants from TdT-01 to **TdT-33**. Coupling efficiencies above 99% are shown in green and display an additional decimal place. Reaction conditions: 2 µM TdT, 1 µM oligo, 5 µM 3’P-dATP, 90s, 60 °C. D) Oligo acceptor sequence bias. Coupling efficiency of TdT-33 with 64 oligo acceptors comprising the full set of 3’-terminal N1N2N3 sequences was measured in a 90-second reaction. Measurements denoted with * indicate a 180 second reaction. Created with BioRender.com.

The stability of the TdT variants improved over the course of evolution. To demonstrate this improvement, residual TdT activities after thermal challenge were assayed using ddGTP as a surrogate substrate, as early variants had minimal measurable activity on 3’P-dNTP substrates. Under these conditions, **TdT-32** retained full activity up to 64°C, a 20°C improvement over the wild-type enzyme (TdT-01). Importantly, these stability gains were not achieved by steady improvements. Over the course of evolution, losses of stability were concomitantly observed with improvements in specific activity, but these losses were recoverable in subsequent rounds. For example, **TdT-22** was less thermostable than **TdT-18** but showed far less oligo acceptor sequence bias. Activity-stability trade-offs such as this underline the importance of continuously screening for the variety of traits needed in the final desired enzyme.

Variants after **TdT-13** gave measurable activity with 3□P-dNTPs and some oligonucleotides under the process-like conditions of short reaction times (90 seconds) and elevated temperatures (60 °C). In assays measuring the activity of six TdT variants (covering **TdT-13** through **TdT-32**) at different reaction temperatures (without pre-incubation), round-over-round improvements with 3’P-dNTP substrates was observed alongside increases in the maximum conversion achieved (Figure 2a). Maximum conversion is also observed at higher reaction temperatures with later variants. Continued improvement of stability alongside activity led to variants that displayed high conversion at temperatures above the 60 °C desired process condition, thereby maintaining a buffer of thermostability for a robust process.

In addition to stability, 3’P-dNTP loading needed to be optimized because reagent costs are a major driver for DNA synthesis cost. While early rounds of evolution were run with 200 molar equivalents of 3’P-dNTP, the molar equivalent excess was reduced 40-fold over consecutive rounds of evolution, with **TdT-32** reaching high conversions at 5 molar equivalents 3’P-dNTPs (Figure 2b). Nearly even incorporation of each 3□P-dNTP was observed with later evolution variants, although pyrimidines remained slightly more favored than purines.

A panel of 16 substrates was screened under target process conditions with 14 TdT variants selected from the evolution lineage (Figure 2c). No activity was observed with early variants under these conditions, but by round 13, TdT variants had high activity on a few substrates, particularly oligos terminating with cytidine, and the main evolutionary pressure became decreasing sequence bias. Activity was observed to increase steadily up to **TdT-22**, at which point there were trade-offs between activity and other traits that were also being targeted for improvement, such as stability and selection against by-products. With **TdT-28**, nearly all sequences reached >90% conversion, and with **TdT-33**, reactions with all but one sequence reached 99% conversion (Figure 2d).

In conclusion, variant **TdT-33** was found to meet most of the key enzyme performance targets, thereby providing proof of concept for a polymerase compatible with an all-enzymatic DNA synthesis platform. Aided by the development of efficient screening protocols and robust, fast analytical methods, **TdT-33** was developed within 14 months of the onset of the evolution campaign. Throughout evolution, TdT’s soluble expression, thermostability, 3’P-dNTP loading, and oligo acceptor sequence bias were maintained or improved. Even after 32 rounds of evolution, improvements in each desired trait continued to be observed, including activity on the most challenging substrate pairs. Additional evolution has been projected to further improve performance, and continued enzyme evolution and process development are ongoing to realize an enzymatic route to long, high-purity synthetic DNA.

## Acknowledgements

We thank David Entwistle, Bill Efcavitch, Deanne Sammond, and Chris Wilson for helpful discussion and collaboration. We thank Ray Blume, Rossana Carillo, Stephanie Hanna, Kathleen Marsters, Steven Navichoque, Sunila Piplani, Kin Lei Seong, Michael Whitehorn, Erika Jane Dumlao, Abigail Bautista, and Dominique Cortez for technical assistance.

## Conflict of Interest

The authors were employees of Codexis, Inc. at the time this work took place.

## Notes

### Competing Interest Statement

The authors declare the following competing financial interest(s): this work has been included in patent application(s) with the United States Patent and Trademark Office and Non-US patent offices. They are as employees and shareholders of Codexis Inc.

